# The influence of carbon dioxide on cerebral metabolism and oxygen consumption: combining multimodal monitoring with dynamic systems modelling

**DOI:** 10.1101/2023.07.24.550309

**Authors:** David Highton, Matthew Caldwell, Ilias Tachtsidis, Clare E Elwell, Martin Smith, Chris E Cooper

**Author notes:** **Correspondence:** Chris Cooper.

## Abstract

Hypercapnia increases cerebral blood flow, but the effect on cerebral metabolism in humans remains incompletely understood. Either increased or reduced oxygen consumption has been predicted from Fick models incorporating cerebral oxygen extraction fraction and cerebral blood flow. Hypercapnia also results in oxidation of cytochrome *c* oxidase, complex IV of the mitochondrial respiratory chain, implicating a change in cellular metabolism. The aim of this study was to combine systems modelling with non-invasive measurements of cerebral tissue oxygenation, cerebral blood flow, and cytochrome *c* oxidase redox state to evaluate any metabolic effects of hypercapnia. Cerebral tissue oxygen saturation and cytochrome oxidase redox state were measured with broadband near infrared spectroscopy and cerebral blood flow velocity using transcranial Doppler ultrasound. Data collected during 5-minutes hypercapnia in human volunteers were analyzed using a Fick model to determine changes in brain oxygen consumption and a mathematical model of cerebral hemodynamics and metabolism (BrainSignals) to inform on the potential mechanisms of any observed changes. Systemic physiology - blood pressure, arterial oxygen saturations and end-tidal carbon dioxide - served as model inputs. The extent of the hypercapnic oxidation of cytochrome oxidase required modifications of the BrainSignals model; simulations compared two possible modifications that could cause this oxidation - a *decrease* in metabolic substrate supply or an *increase* in metabolic demand. Only the decrease in substrate supply was able to explain both the enzyme redox state changes and the Fick calculated drop in brain oxygen consumption. These modelled outputs are consistent with, but do not prove, previous reports of CO_2_ inhibition of succinate dehydrogenase, complex II of the mitochondrial respiratory chain. These findings suggest that hypercapnia may have physiologically significant effects suppressing oxidative metabolism in humans and perturbing mitochondrial signaling pathways in health and disease.

## 1 INTRODUCTION

Despite being studied for over sixty years, there is still a lack of consensus on the effect of arterial carbon dioxide on brain function. The pioneering work of Kety and Schmidt (Kety and Schmidt, 1946) demonstrated effects of CO_2_ on cerebral blood flow (CBF); hypercapnia induces dilation of cerebral arteries and arterioles and increases CBF, whereas hypocapnia causes constriction and decreases CBF. Though the molecular mechanism of this effect is still under debate (Yoon et al., 2012), the control of CBF by CO_2_ is highly reproducible, is not controversial and is viewed as an important system for controlling the cerebral circulation. At the same time as measuring CBF, Kety and Schmidt (Kety and Schmidt, 1946), measured the effect of hyperventilation on the cerebral metabolic rate for oxygen (CMRO_2_). However, subsequent studies, revealed that the relationship between CO_2_ and CMRO_2_ is less well-defined and the underlying mechanisms of it less-well understood. In a review in 1980, entitled “Cerebral Metabolic Rate: a controversy” Siesjö (Siesjo, 1980) noted a range of studies showing an increase, decrease or no effect on CO_2_ on CMRO_2_, concluding “it was disconcerting that 30 years after the first quantitative report we still do not know how hypercarbia (hypercapnia) affects cerebral metabolic rate”.

Fast forwarding thirty more years the controversy is unresolved (Yablonskiy, 2011), with more recent studies reporting a decrease (Xu et al., 2011; Peng et al., 2017), no effect (Chen and Pike, 2010; Jain et al., 2011) or an increase (Jones et al., 2005) in CMRO_2_ with hypercapnia. The effects are usually rather small. For example, one study showed that 5% CO_2_ inhalation led to a 54% increase in CBF, but a more modest 13% decrease in CMRO_2_ (Xu et al., 2011). Measuring CMRO_2_ is inherently more problematic and variable than measuring CBF, if only because the standard method carries all the errors of a CBF measurement, and then adds variability due to measures of arterial and venous oxygen saturation. Even though there are now tools that were unavailable to Kety and Schmidt to measure flow and metabolism, such as MRI and PET, most current measures of CMRO_2_ still share the advantages – and pitfalls – of their steady state method (Kety and Schmidt, 1946). Therefore it is possible, as some have stated explicitly (Yablonskiy, 2011), that the experimental variability observed is wholly due to systematic differences in the measurement techniques used.

Yet, even a small change in this difficult to measure physiological parameter is important. Even a 5% change in CMRO_2_ could have profound physiological and pathophysiological consequences. For example, changes in arterial pCO_2_ influence changes in cerebral blood flow during exercise (Ogoh and Ainslie, 2009), prolonged apnea (Bain et al., 2016) respiratory lung disease (Contou et al., 2015) and in ventilated patients undergoing surgery or in critical care (Mutch et al., 2020). Yet the metabolic consequences of these changes – independent of changes in pO_2_ – are unclear. Thus the resolution of this debate in human subjects holds considerable physiological and clinical importance. Additionally CO_2_ changes are frequently used to calibrate spectroscopic and imaging methods; the assumption is that CO_2_ only mediates flow, with no effect on metabolism (Hoge, 2012). Techniques that assume CO_2_ is isometabolic, such as calibrated fMRI, would therefore be compromised if CO_2_ significantly affected CMRO_2_.

Near Infrared Spectroscopy (NIRS) is a non-invasive technique for measuring tissue hemoglobin oxygen saturation and mitochondrial cytochrome oxidase redox state in vivo (Elwell and Cooper, 2011). NIRS has been suggested as being able to estimate CMRO_2_ by measuring blood flow and the difference between arterial and tissue oxygen saturation (Boas et al., 2003). However, we have been developing an alternative method to measure CMRO_2_, combining broadband NIRS measures of hemoglobin and mitochondrial cytochrome oxidase with a dynamic systems model (BrainSignals) of brain oxygen delivery and metabolism (Banaji et al., 2008).

In this paper we used a previously generated adult volunteer data set to compare direct (Fick) and mathematically modelled (BrainSignals, Figs 1 and 2) NIRS measurements of the effects of CO_2_ on CMRO_2_. Our intention was to optimize our model using the measured changes in the mitochondrial cytochrome oxidase redox state and thus to test whether a combination of a direct mitochondrial measurement with brain systems modelling can both confirm the effect of CO_2_ on brain energy metabolism and inform on any possible underpinning biochemical mechanisms.

**Fig 1.**
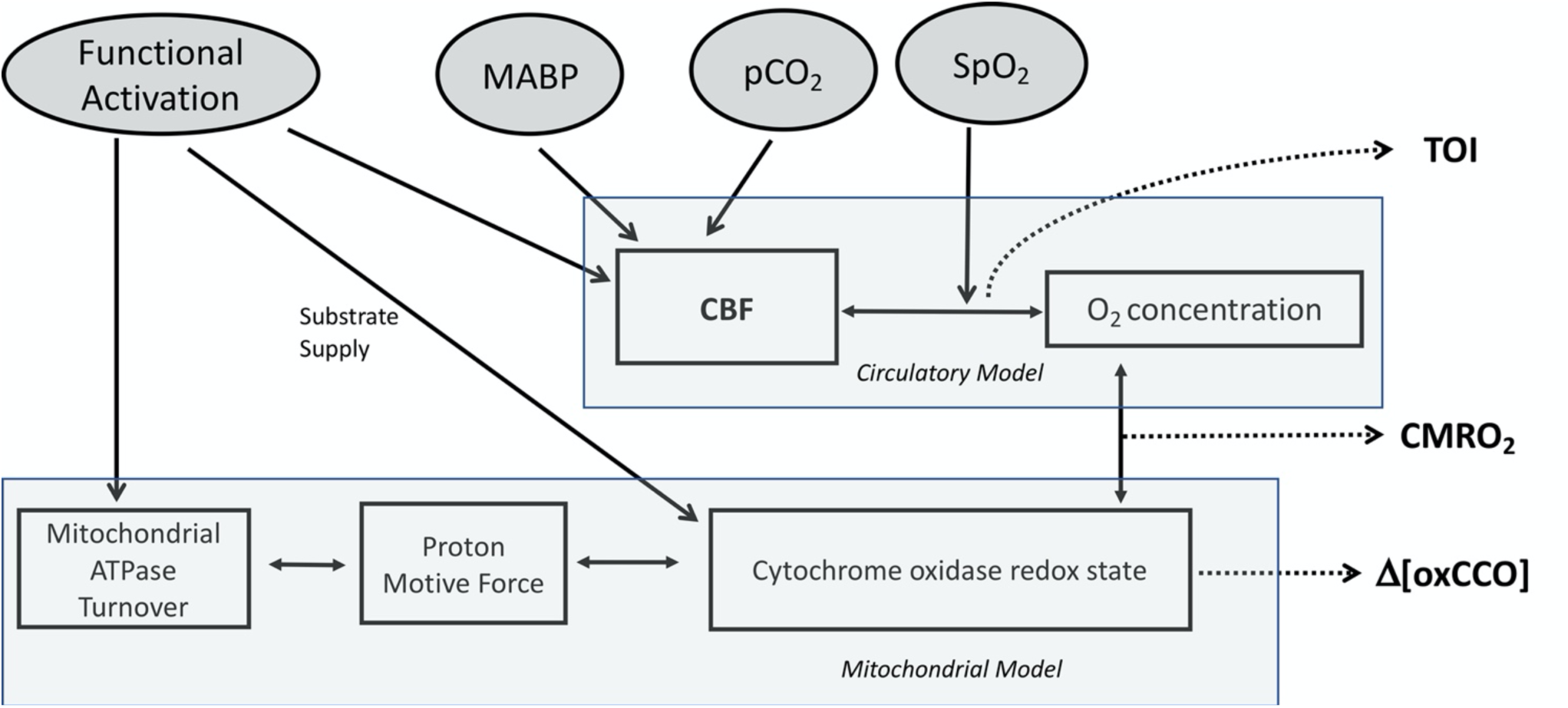
Basic Structure of BrainSignals Model. Figure adapted from Banaji et al (2008) (Banaji et al., 2008). Summary of the main inputs, variables and processes in the model. Circulatory and mitochondrial sub models are linked by the interactions of O2 concentration and cytochrome oxidase redox state. Four physiological inputs feed into the model MABP, pCO2 and SpO2 feed into the circulatory model via effects on CBF (MABP and pCO2) and O2 concentration (SpO2); functional activation feeds into both the circulatory sub model (via effects on CBF) and the mitochondrial sub model via modifying mitochondrial ATPase turnover (ADP/ATP ratio) and the supply of mitochondrial reducing equivalents. Outputs of particular interest in this study are highlighted in bold (**CBF, TOI, CMRO2 and oxCCO**).

**Fig 2.**
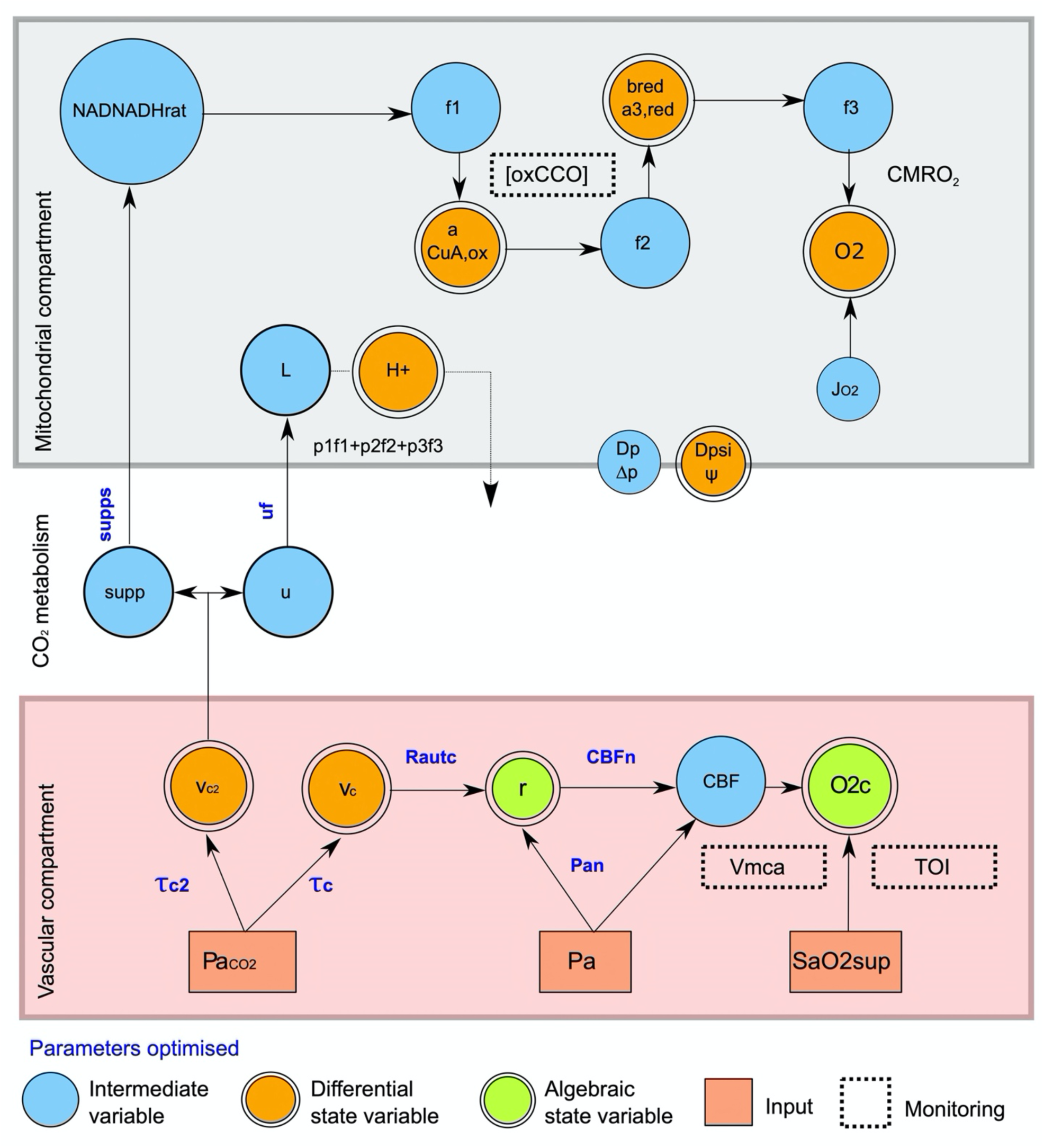
Summary of key dependencies in modified BrainSignals model. Arterial pCO_2_ (Pa_CO2_), mean arterial blood pressure (Pa) and arterial oxygen saturation (SaO2sup) serve as model inputs determining the state variables. [oxCCO] is determined from the simulated CCO CuA center (a), Vmca from CBF, and TOI via oxygen extraction fraction, and the arterial (r) and venous volumes. The hypothesized CO_2_ metabolic effects are simulated via supps (substrate supply) and *u*f (metabolic demand) and their effect on NAD oxidation (NADNADHrat) and proton (H^+^) re-entry into the mitochondrial matrix (L) respectively. **VASCULAR COMPARTMENT** Pa_CO2_ , Arterial pCO_2_; Pa, mean arterial blood pressure; SaO2sup, arterial oxygen saturation; V_c2_, vascular effects of CO_2_ ; τ_c2_time constant filtering vascular effects of CO_2_; V_c_, metabolic effects of CO_2_ ; τ, time constant filtering metabolic effects of CO_2_; Rautc, magnitude of CO2 reactivity on vasculature; r, typical blood vessel radius; Pan, normal arterial blood pressure; CBFn, normal cerebral blood flow; CBF, cerebral blood flow; O2c, capillary O2 concentration; Tissue Oxygenation (TOI); Vmca, Blood velocity in the middle cerebral artery **CO2 metabolism** Supp, Relative supply of reducing substrate to mitochondria; supps, influence of CO_2_ on supply of reducing equivalents; u, normal metabolic demand ; uf, influence of CO_2_ on metabolic demand. **MITOCHONDRIAL COMPARTMENT** NADNADHrat, mitochondrial NAD/NADH ratio; Relative supply of reducing substrate to mitochondria; L, rate of proton return to the mitochondrial matrix; H+, mitochondrial proton concentration; f1, reduction rate for the CuA center of cytochrome oxidase by NADH (essentially rate of electron transfer via complex I and III of electron transfer chain); f2 reaction rate for the reduction of cytochrome oxidase haem a3 by electrons from CuA; f3 Reaction rate for the reduction of O_2_ by haem a3; pf1+pf2+pf3, protons pumped out of mitochondria by the three parts of the mitochondrial electron transfer chain modelled by f1,f2 and f3; CuAox, concentration of oxidized cytochrome c oxidase CuA center; a3r red; concentration of reduced cytochrome a3, O2, mitochondrial oxygen concentration; oxCCO, broadband NIRS measured CuA signal; CMRO2, cerebral metabolic rate for oxygen; J_O2_ , oxygen flux from blood to tissue; Dp Δp, proton motive force across the mitochondrial inner membrane; Dpsi Δψ, membrane potential gradient across the mitochondrial inner membrane.

## 2 MATERIALS AND METHODS

### 2.1 The BrainSignals Model

In 2005 our group developed a complex model to investigate brain circulation and metabolism, called BrainCirc (Banaji et al., 2005). This model incorporated equations representing blood flow, ion channel activity in the vascular smooth muscle and respiration from glycolysis to the electron transport chain. The circulatory portion of the model was derived from Ursino and Lodi (Ursino and Lodi, 1998). In 2008, we further developed the model to allow improved optimisation against experimental data, in particular non-invasive NIRS-derived measurements of changes in hemoglobin oxygenation and the mitochondrial cytochrome oxidase redox state. Model complexity was minimised by removing or simplifying physiological components regarded as nonessential to the basic observed behaviours. This simplification resulted in a new model called BrainSignals (Banaji et al., 2008). The BrainSignals model consists of two submodels: a simplified version of the Ursino and Lodi model of the cerebral circulation (Ursino and Lodi, 1998), and a model of mitochondrial metabolism related to those presented by Korzeniewski (Korzeniewski and Zoladz, 2001) and Beard (Beard, 2005). The two submodels are linked via the processes of oxygen delivery and consumption. Although BrainSignals was intentionally simpler and more comprehensible than BrainCirc, it modelled the electron transport chain in more detail in order to simulate the NIRS-derived ΔoxCCO signal. BrainSignals has been validated and its outputs compared with measurements during hypoxia challenges (Jelfs et al., 2012), hypercapnia challenges (Moroz et al., 2012b) and anagram solving tasks in healthy adults (Kolyva et al., 2012). Adaptations of the BrainSignals model framework outlined in (Banaji et al., 2008) have been made to allow for different inputs and outputs, or to test specific experimental hypotheses. These included simulating glycolysis, lactate dynamics, the tricarboxylic acid (TCA) cycle and hypoxia in the neonatal piglet brain (Moroz et al., 2012a), simulating preclinical models of hypoxic-ischemic brain damage (Caldwell et al., 2015), and simulating cerebrovascular pressure reactivity in critically brain-injured patients (Highton et al., 2013).

The BrainSignals model is described in detail in Banaji et al (Banaji et al., 2008). The modifications of the BrainSignals model used in this paper are described in detail in Fig S1 and are available for download at https://github.com/bcmd/co2.

### 2.2 Experimental study

Data from a previous study investigating Δ[oxCCO] during manipulation of cerebral oxygen delivery in healthy volunteers were analyzed as part of the current study. The previous study, which was approved by the Research Ethics Committee of the National Hospital for Neurology and Neurosurgery and Institute of Neurology (04/Q0512/67) and conducted in accordance with the declaration of Helsinki, is reported elsewhere (Kolyva et al., 2014). The present work specifically focuses on one paradigm of that original study. In brief, following written informed consent, hypercapnia was induced in 15 healthy adult volunteers by the addition of 6% CO2 to the inspired gas mix for 300 seconds, targeting an increase of ∼2 kPa in end-tidal partial pressure of CO2. Monitoring during the period of hypercapnia included: continuous non-invasive arterial blood pressure (PortaPres, Finapres Medical Systems, Netherlands), pulse oximetry (Oxypleth, NovaMetrix, MA), end-tidal CO_2_ (IntelliVue MP50, Philips, Netherlands), transcranial Doppler (TCD) ultrasound of the middle cerebral artery (DWL DopplerBox, Singen Germany), and a hybrid optical spectrometer (HOS) combining frequency and broadband NIRS (Ghosh et al., 2012; Kolyva et al., 2012). The NIRS optodes were located ipsilateral to the TCD recording. Systemic signals (mean arterial blood pressure (MABP), arterial oxygen saturation (SpO_2_), end-tidal CO_2_ (E_T_CO_2_)) and TCD were gathered online at 100Hz and saved for offline analysis. HOS recordings consisted of broadband intensity at 4 source detector separations (20mm, 25mm, 30mm, 35mm) and frequency domain recordings (30mm, 35mm) using an ISS Oximeter, model 96208 (ISS Inc, Champaign, IL, USA). The systems were time multiplexed over the same region alternating every 1.5sec.

Broadband spectroscopy (780nm-900nm) was used to derive [oxCCO] changes as previously described using an established technique and the UCLn algorithm (Kolyva et al., 2014). Individual recordings were pathlength corrected using the differential pathlength factor derived from frequency domain spectroscopy. Cerebral TOS, using the tissue oxygenation index (TOI), was calculated from the slope of attenuation (20mm-35mm) in the broadband data (740-900nm) according to equation (1). TOI may be particularly sensitive to cerebral water concentration and the wavelength dependence of light scattering (h eqn 1.). Because the CMRO_2_ calculation is likely to be highly susceptible to small errors in TOI, additional measures were taken: 1) fitting the spectra for HbO_2_, HHb and H_2_O, 2) adjusting the value of h (eqn 1) to the measurements obtained from the frequency domain group data.

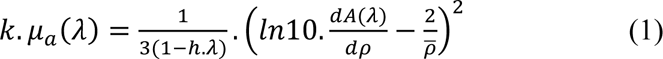

The attenuation coefficient μa multiplied by a constant *k* at a specific wavelength *λ* is determined via eqn. (1). Where *h* is the wavelength dependence of light scattering, *A* light attenuation and *ρ* source detector separation.

MABP was time integrated using an automated peak detection and cleaning algorithm (Zong et al., 2003).Signals were then low pass filtered (0.25Hz, 5^th^ order Butterworth) and resampled at 1Hz to produce time synchronized data for the modelling process: MABP, E_T_CO_2_, SpO_2_, TCD, TOI and [oxCCO]. The study demonstrated a statistically significant increase in [oxCCO], TOI and Vmca following a hypercapnia challenge (Kolyva et al., 2014). The raw data and model used for the modelling process is downloadable from https://github.com/bcmd/co2.

### 2.3 The modelling process

BrainSignals simulates cerebral oxygen physiology across several compartmentalized physiological scales, using experimental recordings of mean arterial blood pressure (MABP), arterial saturation (SpO_2_) end-tidal CO_2_ (ETCO_2_) and brain activation (u) to predict the behavior of a variety of physiological and experimental parameters, including cerebral blood flow (CBF), brain tissue oxygen saturation (TOS) and mitochondrial cytochrome oxidase redox state ([oxCCO]). Fig 1 illustrates the basic structure of the BrainSignals model, and the experimental model inputs and outputs. Changes in the inputs with time (the experimental measurements in Fig 3) define changes in the outputs (the modelled “measurements” in Fig 4). Model outputs can be directly compared to experimentally defined data (again see Fig 4). Model parameters can then be adjusted to optimize the fit between experimental and modelled parameters, and to inform on the underlying biochemical and/or physiological changes that are occurring, including the derivation of parameters that are not easy to measure using conventional non-invasive techniques, such as CMRO_2_ (Figs 5-7 and Tables 1 and 2).

**Fig 3.**
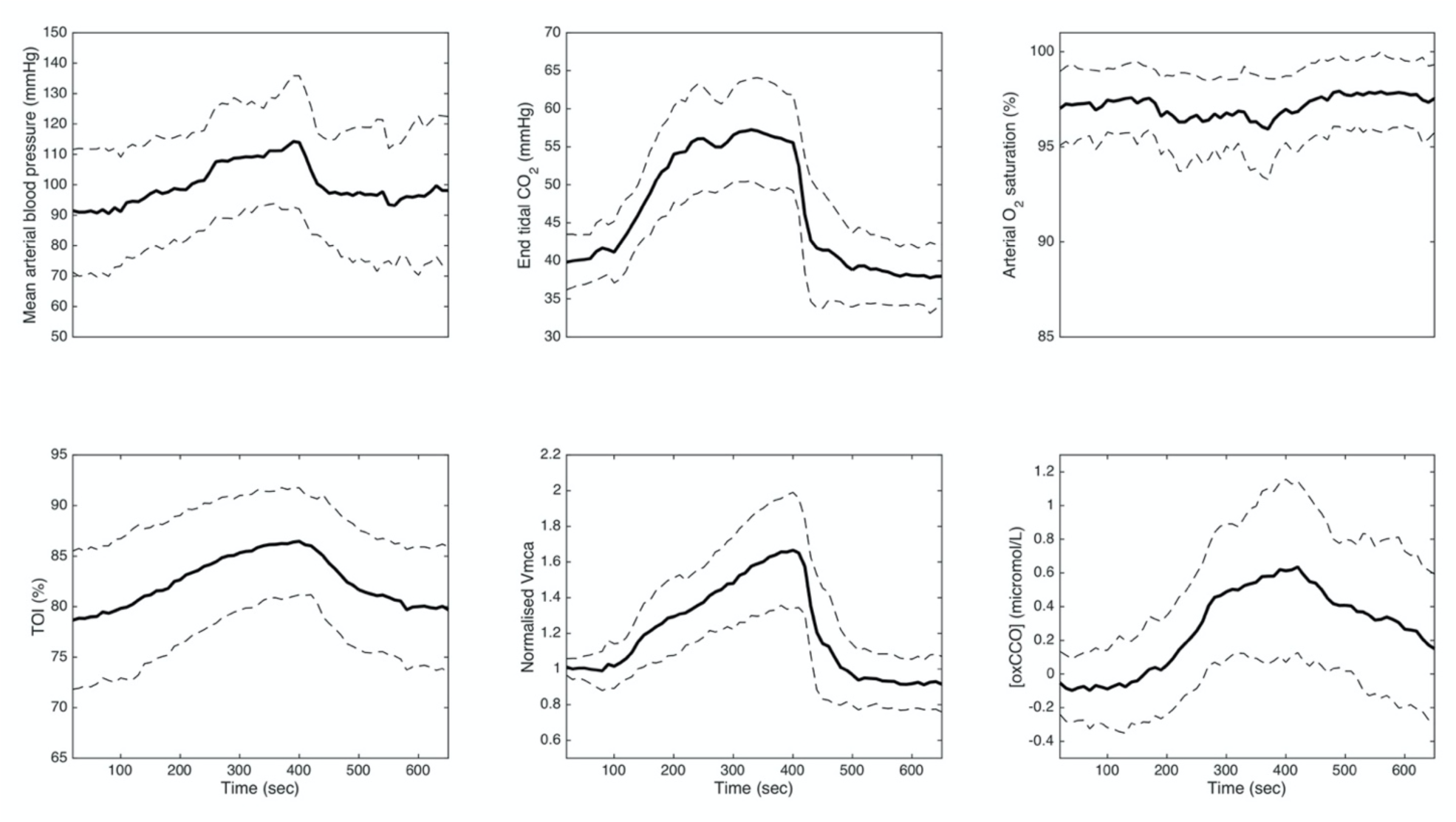
Systemic physiological and cerebral measurements in volunteers. Mean value is in bold (n= 15). Standard deviation is denoted by the dotted line.

**Fig 4.**
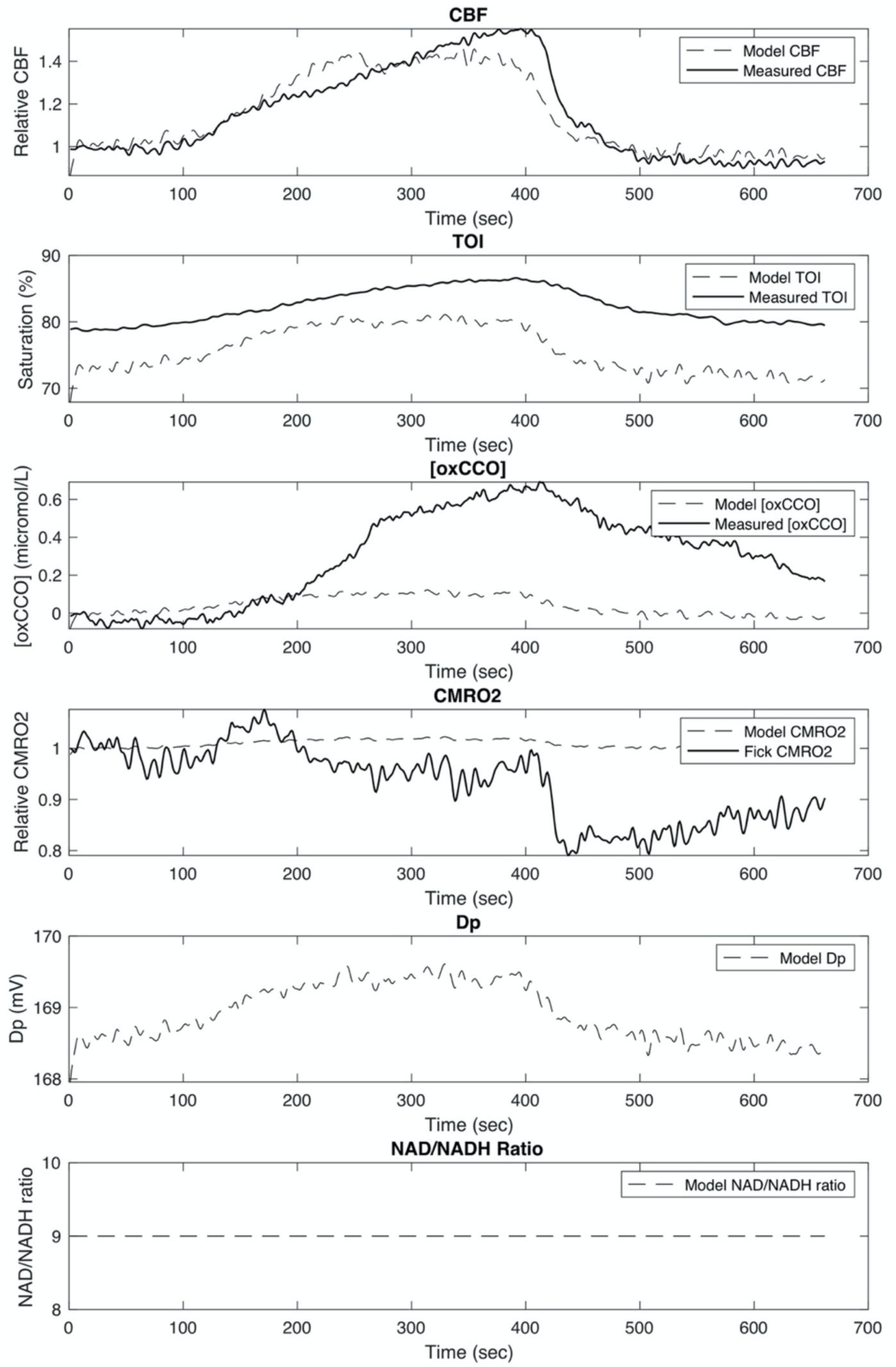
Unoptimized BrainSignals model outputs. The basic BrainSignals model was run using the physiological inputs from Fig 3. Illustrated are the time courses of model outputs relating to cerebral physiology: CBF, CMRO2; biochemistry: Dp (mitochondrial Δp), NAD/NADH (mitochondrial NAD/NADH ratio): measured optical changes (TOI, oxCCO).

**Fig 5.**
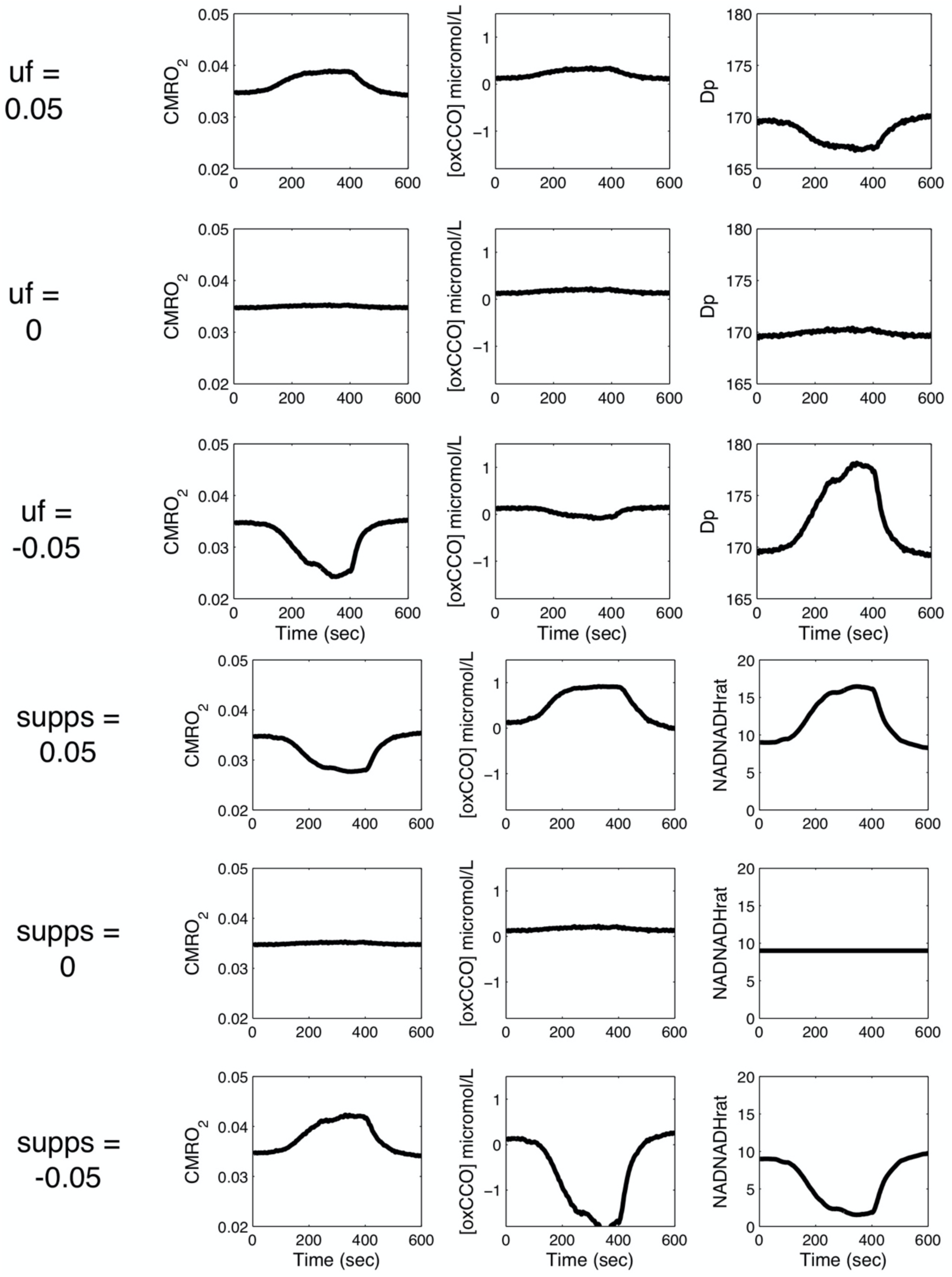
Effect of varying CO_2_ effects on substrate supply and metabolic demand. Selected outputs from modified BrainSignals model using measured inputs from Fig 3 and varying the effects of CO_2_ on ATP demand (uf) or substrate supply (supps). Charts illustrate the effects on CMRO_2_, oxCCO and the dominant parameter affected by substrate supply (mitochondrial NAD/NADH ratio) and ATP turnover (mitochondrial Dp or Δp).

### 2.4 Changes to BrainSignals model

The effect of CO_2_ on the vasculature remained essentially unchanged from the original model. Vascular effects are delayed by a time constant τ_c_ and a first order filter (eqn. 2), as the vascular effects of CO_2_ are typically delayed in the region of 30s (Ursino and Lodi, 1998). Likewise a theoretical CO_2_ effect on metabolism may be delayed on an independent timescale from the vascular effects, and thus an identical but independent first order filter was defined for the metabolic effects of CO_2_ with a time constant (τ_c2_)

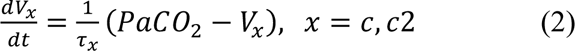

In default model behavior (Fig 1) u is coupled to CBF, simulating the increase in CBF following functional activation of the brain. However, having CBF determined by both CO_2_ and metabolic demand (u) creates unwanted complexity for the present simulation. Therefore any direct effect of u on CBF was disabled by disabling the relevant control parameter (R_u_ = 0). The revised model (Fig 2) therefore assumes that flow changes result from CO_2_ alone, allowing the model to independently control CBF and metabolism.

Variations in modelled parameters were enabled by creating “supply” (supps) and “demand” (uf) parameters that were sensitive to changes in CO_2_ and modified the otherwise fixed model variables of NAD:NADH_rat_ and u. The following parameter regimes were used to evaluate the two proposed mechanisms (1) modifying substrate supply, [**supp_s_,** R_autc_, τ_c,_ τ_c2_, P_an_, CBF_n_] (2) modifying metabolic “demand” [**uf**, R_autc_, τ_c,_ τ_c2_, P_an_, CBF_n_]. Fig 2 illustrates the basic structure of the effects of CO2 on brain blood flow and oxygen consumption and the changes made to the BrainSignals model to enable different effects of hypercapnia on CMRO_2_. This is described in more detail below.

Metabolic substrate supply is currently modelled via a parameter representing the NAD:NADH ratio (NAD:NADH_rat_). A “substrate supply” parameter *supps* inducing an effect of CO_2_ was added (eqn. 3). This simulates an effect of arterial CO_2_ varying electron entry into the electron transport chain proximal to CCO, consistent with variations of NADH demonstrated experimentally following hypercapnia (Gyulai et al., 1982). The term *u*^2*DNADH*^ is used in the original BrainSignals model to change the rate of production of mitochondrial NADH via glycolysis and/or the citric acid cycle. This was modified in our new model via the in introduction of a CO_2_-dependent supp term which allows for positive or negative effects of CO_2_ on substrate supply (eqn. 4).

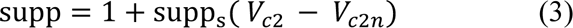

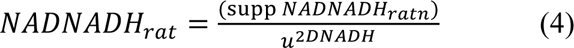

A “metabolic demand” parameter *uf* was added to mediate CO_2_ effects on cerebral metabolic demand (eqn. 5).

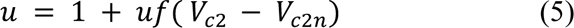

Eqn. 6 and 7 together define the main effects of *u* on cerebral metabolic demand and mitochondrial ATP turnover. L_CV_ defines the rate at which protons re-enter the mitochondrial matrix due to ADP phosphorylation and θ represents the driving force for ATP synthesis. *r*_CV_ controls the scale of L_C_ changes, Δp_CV0_ identifies the Δ*p* where *L_cv_* becomes 0, and Z is a constant. Varying *u*f allows for CO_2_ to increase or decrease the effects of metabolic demand on the mitochondrial proton motive force and hence oxygen consumption,

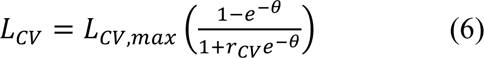

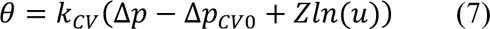

### 2.5 Model analysis

Model inputs included MABP, arterial hemoglobin oxygen saturation and ETCO_2_ (estimation of arterial pCO_2_) were sampled at 1Hz and then converted to group mean values for modelling input. These input data via the model generate simulated outputs for the experimentally measured data: [oxCCO], normalized change in CBF and TOI. Functional Activation (*u)* was the only input in the original BrainSignals model not measured in this study. It was initially assumed that CO_2_ had no effect on brain activity so this model parameter (*u)* was not varied from its default model value of 1. However, to test whether varying pCO2 might have metabolic consequences the effects of (*u)* on mitochondrial substrate supply (via changes in the NAD/NADH ratio) and ATP turnover (via effects on the proton leak) were modelled by introducing two new variables (supps and *u*f respectively, see Fig 2 and previous discussion).

The simulations are controlled in part by model parameters which define key aspects of physiology. Thus, by using a mathematical optimization technique (see later) optimal parameter values can be identified which produce similar modelled and measured data and identify the physiological state. This process is well recognized, but - due to the complexity of such models - it is not possible to individually identify *all* parameters. In a previous sensitivity analysis we identified a number of the key parameters that can induce changes in the cerebral cytochrome oxidase redox state (Russell-Buckland et al., 2019). However, some of these (such as total mitochondrial content) are not relevant to the acute changes in brain physiology that could be perturbed by a change in CO_2_. Therefore, we selected a subset of model parameters related to our hypothesis based on physiological relevance and which specifically addressed our metabolic hypothesis, drawing on previous CO_2_ modelling studies to describe its essential effects on CBF (Ursino et al., 2000a; Ursino et al., 2000b; Moroz et al., 2012b). To investigate the metabolic effect of CO_2_ we used the new model parameters *uf* (metabolic demand) and *supp_s_* (substrate supply), modified by a time constant τ_c2_ specifically designed to describe the metabolic hypotheses outlined in the Results section. Previous CO_2_ mathematical modelling has highlighted the importance of the temporal time course, autoregulation and CO_2_ reactivity to describe CBF (Ursino et al., 2000a; Ursino et al., 2000b; Moroz et al., 2012b). Parameters required to explain these key physiological effects of CO_2_ were selected based on their known physiological variation and previous parameter choices in related models describing CO_2_ reactivity. These are the magnitude of the CO_2_ effect on CBF (*R_autc_*), the temporal delay of this CO_2_ effect (τ_c_), its effect on cerebral autoregulation (P_an_) and the variation of resting CBF (*CBF_n_*). The following parameter regimes were used to evaluate the two proposed mechanisms of CO_2_ affecting CMRO_2_ differing only in the metabolic parameters (supp_s_, uf): 1) modifying substrate supply, [**supp_s_,** R_autc_, τ_c,_ τ_c2_, P_an_, CBF_n_], and, 2) modifying metabolic “demand” [**uf**, R_autc_, τ_c,_ τ_c2_, P_an_, CBF_n_]. Global optimization was performed to fit the parameters using a genetic algorithm (ga, Matlab). This sought to minimize a cost function equally weighted for ΔCBF%, TOI and [oxCCO], when optimally fitted as previously described in Jelfs et al (Jelfs et al., 2012) (eqn. 8), where d(R_x_) represents the distance between measured and modelled variables (x = ΔCBF%, TOI, [oxCCO]). This distance is calculated as the mean absolute difference between measured and modelled variables. Because of differing variable scales γ_R_, a weighting factor is applied (γ_D/012%_ = 0.025 , γ_D456_ = 0.42, γ_D[8!--5]_ = 0.04). This process was repeated 200 times for each optimization, the standard deviation of the parameter variation between these trials is reported as a measure of success for optimization.

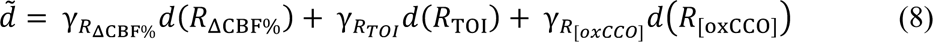

The changes in CMRO_2_ reported from the BrainSignals model post-optimization were compared with a direct calculation from measured ΔSpO_2_, ΔTOI and ΔCBF using a modified Fick model (Boas et al., 2003; Roche-Labarbe et al., 2012) (eqn. 9). This model incorporates the CBF change and the change in the difference in arterial and venous oxygen saturation (the latter calculated from the TOI).

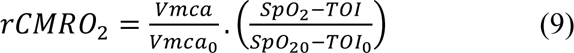

## 3 RESULTS

### 3.1 Experimental study and derived CMRO2

The mean group data are displayed in Fig 3. Hypercapnia results in increased MABP, increases in CBF (Vmca) increases in TOI and increases in [oxCCO], but no change in SpO_2_.

Fig 4 shows the unoptimized BrainSignals output using the physiological inputs from Fig 3 (MABP, SpO2 and ETCO_2_). It can be seen that CBF is quite well modelled, as are changes in TOI (but not the absolute value). The measured increase in oxidation of the cytochrome oxidase CuA center is present, but the magnitude is significantly underestimated in the model. CMRO_2_ was not measured directly, but Fig 4 compares the BrainSignals estimate with that using ΔSpO_2_, ΔTOI and ΔCBF to calculate ΔCMRO_2_ from the modified Fick equation (equation 9). The Fick calculated ΔCMRO_2_ shows a small fall in CMRO_2_ during hypercapnia, as the increase in TOI more than offsets the increase in CBF. In contrast the unmodified BrainSignals model shows a small increase in ΔCMRO_2_ during hypercapnia. Of relevance to later modelling the unmodified BrainSignals shows a very small increase in mitochondrial proton motive force (likely driven by the CBF increase) but no change in the fixed value of NAD/NADH ratio.

### 3.2 Rationale for the modifications of the BrainSignals model to fit the experimental data

The BrainSignals model outputs simulated values of ΔCMRO_2_, ΔTOI, ΔCBF and Δ[oxCCO] based on changes in the input systemic variables only. It does not *a priori* account for the measured NIRS outputs and/or the Fick calculated CMRO_2_. The model was designed to be optimized against experimental data, allowing for changes in the starting system variables. However, in the case of hypercapnia the only relationship between the ETCO_2_ input and the measured NIRS variables in the current model is via changes in CBF (Fig 1). Thus an *increase* in CBF induced by hypercapnia can only increase oxygen delivery and therefore only cause an *increase* in CMRO_2_ (Fig 4), the extent of which will depend on how much, if at all, CBF currently limits CMRO_2._ Therefore to simulate the fall in CMRO_2_ - as indicated by the Fick-derived calculations - structural changes need to be applied to the model. The success of these changes can then be tested by how well they fit the measured NIRS variables as well as the fall in CMRO_2_. Different structural changes can be modelled to cause a hypercapnia-induced decrease in CMRO_2_; the [oxCCO] measure is likely to be the most useful mechanistic discriminator between models as it can report on how mitochondrial oxygen consumption has been altered (Cooper et al., 1994; Banaji, 2006).

Fig 2 illustrates the factors affecting mitochondrial oxygen consumption, how they are currently modelled in BrainSignals and the proposed changes to enable a decrease in CMRO_2_ in hypercapnia. As noted earlier, the effect of hypercapnia in increasing the rate of oxygen delivery to mitochondria can only increase, not decrease oxygen consumption. At the level of mitochondrial cytochrome oxidase oxygen consumption is varied by the rate of substrate delivery to the enzyme (oxygen, protons and reducing agents from the respiratory chain) and the size of the mitochondrial proton electrochemical potential (Δp), comprising the mitochondrial membrane potential (Δψ) and the ΔpH across the inner mitochondrial membrane (Brand and Murphy, 1987). The next section will deal in more detail with the three substrates (O_2,_ protons, and mitochondrial reducing equivalents) that can increase cytochrome oxidase activity and the one product (Δp) that can decrease it.

### 3.3 Modifying cytochrome oxidase activity in BrainSignals

#### 3.3.1 Oxygen

As the oxygen delivery part of the BrainSignals has been shown to function well in simulating a range of experimental data, including hypercapnia induced blood flow changes, (Fig 4), it was also decided not to modify this part of the model.

#### 3.3.2 Protons (pH)

Unlike some modified versions (Moroz et al., 2012a), the basic BrainSignals model does not allow for mitochondrial pH changes, independent of changes in Δp. Experimental evidence suggests that even 1h of hypercapnia only causes a 0.1 fall in brain pH (Nishimura et al., 1989). The effect on cytochrome oxidase activity of such a small change is negligible and, if anything, increases oxygen consumption (Thornstrom et al., 1988), rather than the decrease we see using the Fick equation (Fig 4). pH *changes* across the inner mitochondrial membrane (as opposed to the absolute pH value) are, however, important and already factored into the model as a component of Δp. Therefore, it was decided not to increase the complexity of our changes by explicitly including changes in intramitochondrial pH into the revised model.

#### 3.3.3 Mitochondrial reducing equivalents

The rate of oxygen consumption is a function of the driving force (redox potential) of cytochrome *c* which acts to increase consumption (Murphy and Brand, 1987). The driving force for cytochrome *c* reduction can be changed in our model by varying the rate of substrate (electron) delivery, effectively portrayed in the basic model by a fixed NAD/NADH redox state NAD:NADH_rat_ . Decreasing the redox driving force (by increasing NAD:NADH_rat_) will decrease oxygen consumption.

#### 3.3.4 *Δp*

Increasing Δp will decease oxygen consumption (Murphy and Brand, 1987). The size of Δp can be altered by the rate of ATP turnover, effectively portrayed in the model by the demand parameter u. Decreasing u will increase Δp and hence decrease oxygen consumption.

It was therefore decided to modify BrainSignals to accommodate CO_2_-induced changes in the rate of delivery of mitochondrial reducing equivalents to cytochrome oxidase (via the NAD/NADH ratio) and the size of Δp (via the demand parameter u.

### 3.4 Details of the BrainSignals modifications

In the original model metabolic demand was simulated using a parameter *u* originally designed to simulate a variety of distinct effects of functional activation (changes in ADP/ATP ratio, mitochondrial substrate supply and blood flow). The parameter *u* therefore has three affects in BrainSignals, one vascular and two metabolic; it increases CBF, increases substrate supply (via glycolysis/ τ_c_ cycle effects on the NAD/NADH ratio) and increases proton entry to the mitochondrial matrix L_CV_ (via changes in metabolic ATP demand).

In the original model, pCO_2_ can only affect mitochondrial metabolism by effects on CBF and consequent changes in mitochondrial O_2_ concentration. As this proved inadequate to explain the experimental behavior additional CO_2_ effects on mitochondria were enabled by allowing CO_2_-driven changes in substrate supply or ATP demand. These two distinct metabolic consequences were modelled by introducing two new variables (supps and *u*f respectively, see Fig 2 and Materials and Methods for more details). A CO_2_-induced decrease in metabolic demand is modelled by a negative uf and an increase by a positive uf. A CO_2_-induced decrease in substrate supply is modelled by a positive supps (as supps increases the NAD/NADH ratio) and an increase in substrate supply by a negative supps.

### 3.5 Comparison of different methods of simulating hypercapnia-induced CMRO_2_ changes

Fig 5 illustrates the effect of the different changes in modelling on the BrainSignals outputs during the hypercapnic challenge illustrated in Fig 3. A CO_2_-induced decrease in metabolic demand (uf - 0.05) decreases oxygen consumption via an increase in the mitochondrial proton electrochemical potential (Δp), whereas a CO_2_-induced decrease in substrate supply (supps +0.05) decreases oxygen consumption via an increase in the mitochondrial NAD/NADH ratio. In principle, measures of Δp and NAD/NADH could therefore be used to discriminate the mechanism of any observed reduction in CMRO_2_. However, these measures cannot be performed noninvasively in the adult human brain. However, both Δp and NAD/NADH influence the redox state of the cytochrome oxide Cu_A_ center. This can be measured (via the oxCCO NIRS signal) and responds differently to changes in metabolic supply or demand. Decreasing CMRO_2_ via a decrease in demand (uf = −0.05) increases Δp and decreases oxCCO, whereas decreasing CMRO_2_ via a decrease in supply (supps = 0.05) increases NAD/NADH and increases oxCCO.

Demand and substrate supply changes are modelled in Fig 6 and Fig 7 respectively, with parameter optimization against the experimental neuromonitoring data (Table 1a and b). To differentiate the model sensitivity to oxygen extraction and oxCCO measurements three optimizations sets are displayed. These examine changes in: cellular oxygen metabolism ([oxCCO] alone); tissue oxygen delivery and extraction (TOI and CBF alone); and oxygen delivery, metabolism and extraction (CBF, TOI and [oxCCO]).

**Fig 6.**
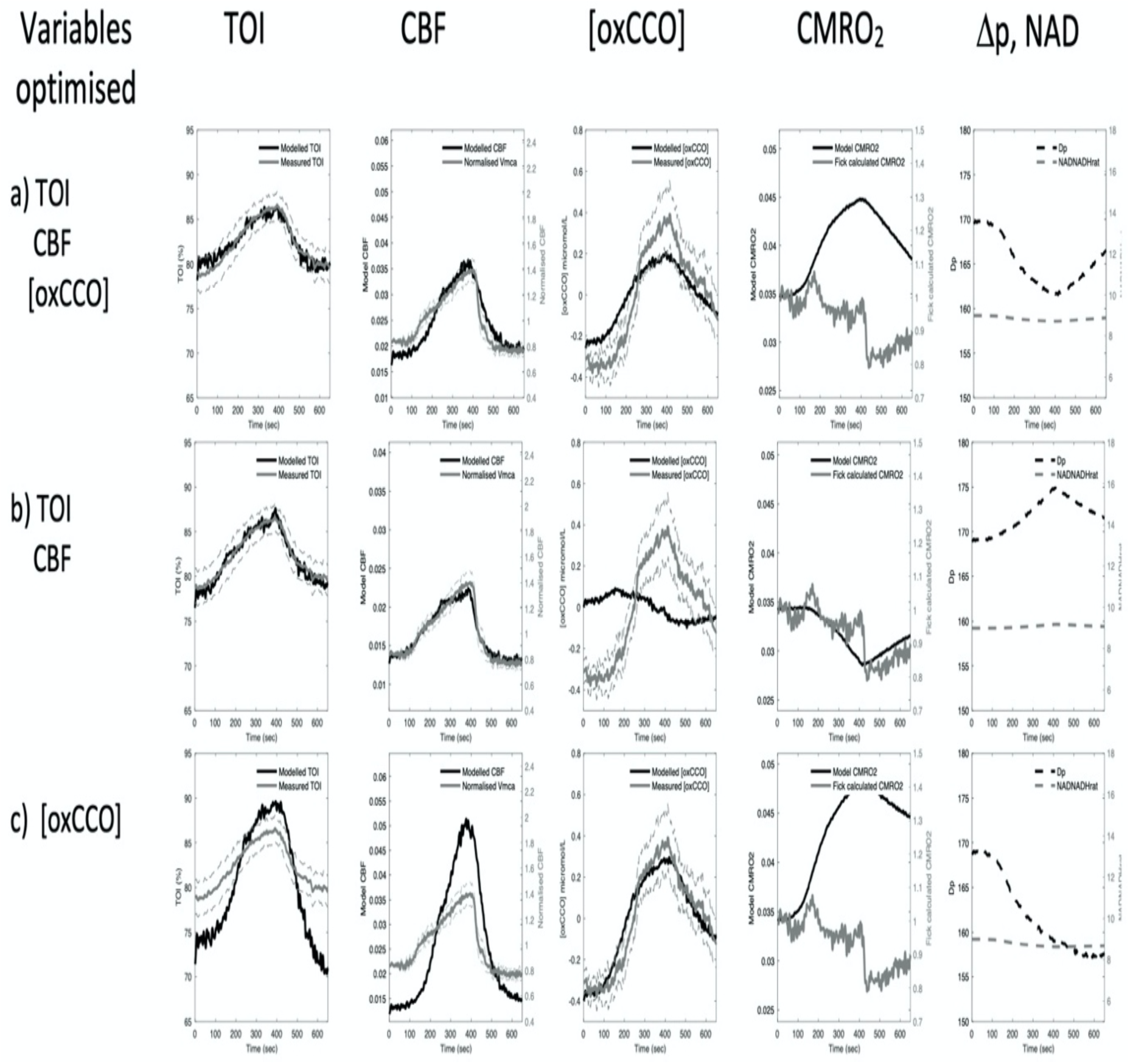
Demand optimization. Model simulation of a change in metabolic demand (parameter uf). In each row different monitoring variables are used to achieve the parameter optimization. It can be seen that no optimization fits the data well (large difference between measured and modelled data), and this produces predictions of both increased and decreased CMRO_2_. Full parameter values are displayed in Table 1a Dashed grey lines denote the standard error of the means of the measured data. For abbreviations see Figure 5 legend.

**Fig 7.**
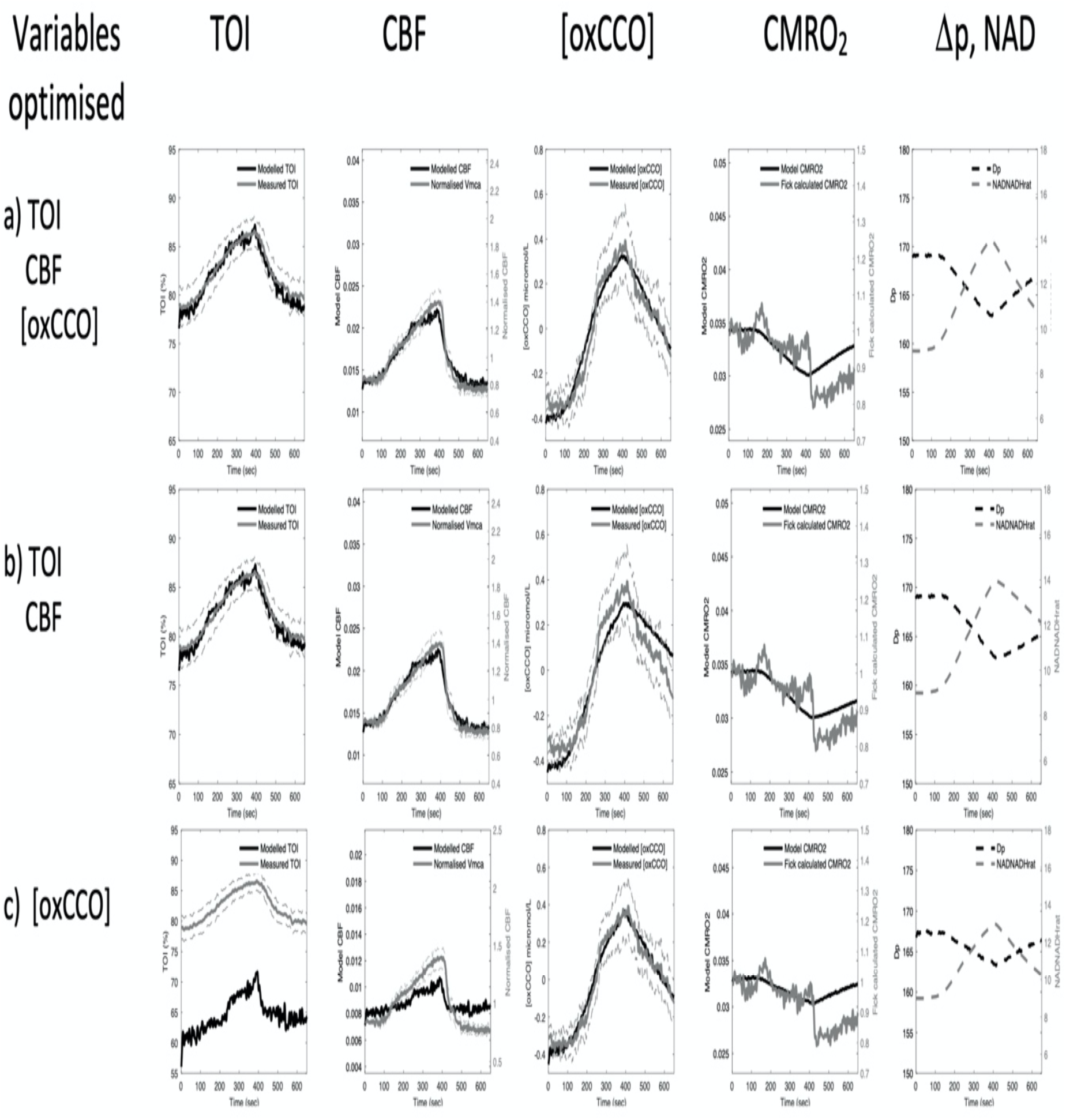
Supply optimization. Model simulation of a change in substrate supply (parameter supp_s_). In each row different monitoring variables are used to achieve the parameter optimization. It can be seen that there is generally a good fit of modelled and measured data and this produces predictions of a small decrease in CMRO_2_. Full parameter values are displayed in Table 1b. Grey dashed lines denote the standard error of the means of the measured data. For abbreviations see Figure 5 legend.

**Table 1a.** Optimization results from demand optimization.

| *Parameters* | *Basal value* | *[oxCCO]*<br>*optimization* | *TOI/CBF*<br>*optimization* | *TOI/CBF/[oxCCO]*<br>*optimization* |
| --- | --- | --- | --- | --- |
| **Demand** | uf (0) | 1.69 ± 0.09 | -0.10 ± 0.03 | 0.33 ± 0.12 |
| **CO<sub>2</sub> reactivity** | R<sub>autc</sub> (2.2) | 9.99 ± 0.25 | 4.66 ± 1.60 | 4.88 ± 0.70 |
| **Time constant CO<sub>2</sub>** | τ<sub>c</sub> (15 sec) | 99.96 ± 0.33 | 24.60 ± 11.34 | 78.24 ± 4.68 |
| **Time constant**<br>**metabolic effects CO<sub>2</sub>** | τ<sub>c2</sub> (15 sec) | 789.32 ± 29.49 | 510.79 ± 142.79 | 188.08 ± 24.88 |
| **Autoregulation** | P<sub>an</sub> (100mmHg) | 68.65 ± 1.73 | 143.03 ± 19.6 | 73.55 ± 18.25 |
| **CBF constant** | CBF<sub>n</sub> (0.0125 ml/ml<br>tissue/sec) | 0.0096 ± 0.0005 | 0.00225 ± 0.00220 | 0.00179 ± 0.0052 |
| **Error** |  |  |  |  |
| **TOI** | (%) | 3.87% | 0.45% | 0.56% |
| **CBF** | (%) | 27.78% | 2.99% | 7.66% |
| **[oxCCO]** | (micromol/L) | 0.00510 | 0.244 | 0.105 |

**Table 1b.** Optimization results from substrate supply optimization.

| *Parameters* | *Basal value* | *[oxCCO]*<br>*optimization* | *TOI/CBF*<br>*optimization* | *TOI/CBF/[oxCCO]*<br>*optimization* |
| --- | --- | --- | --- | --- |
| **Substrate supply** | supp<sub>s</sub> (0) | 0.0424 ± 0.0024 | 0.0985 ± 0.0085 | 0.0565 ± 0.0034 |
| **CO<sub>2</sub> reactivity** | R<sub>autc</sub> (2.2) | 0.01 ± 0.17 | 4.67 ± 1.32 | 4.55 ± 1.23 |
| **Time constant CO<sub>2</sub>** | τ<sub>c</sub> (15 sec) | 92.83 ± 18.55 | 26.20 ± 6.39 | 26.96 ± 5.30 |
| **Time constant metabolic effects CO<sub>2</sub>** | τ<sub>c2</sub> (15 sec) | 243.00 ± 5.85 | 605.98 ± 76.32 | 273.03 ± 20.89 |
| **Autoregulation** | P<sub>an</sub> (100mmHg) | 67.57 ± 5.15 | 142.75 ± 13.12 | 144.86 ± 6.45 |
| **CBF constant** | CBF<sub>n</sub> (0.0125 ml/ml tissue/sec) | 0.0011 ± 0.0012 | 0.00226 ± 0.0012 | 0.0028 ± 0.0009 |
| **Error** |  |  |  |  |
| **TOI** | (%) | 17.70% | 0.46% | 0.49% |
| **CBF** | (%) | 149% | 3.03% | 3.43% |
| **[oxCCO]** | (micromol/L) | 0.0043 | 0.0071 | 0.0040 |

Simulated changes in metabolic demand (uf) poorly replicate the experimental measurements (Fig 6, Table 1a). Optimizing against the [oxCCO] variable alone can model the measured increase in [oxCCO] as an increase in demand, decreases (Δp) and oxidizes the Cu_A_ center in the enzyme (Fig 6c). However, although a TOI and CBF increase during hypercapnia is modelled, the optimization overestimates the magnitude of the experimental changes. As a consequence, the model optimized CMRO_2_ shows a significant hypercapnic increase in CMRO_2_, differing in both sign and magnitude from the Fick-derived CMRO_2_. Optimizing against TOI and CBF alone (Fig 6b) shows good fits to both the experimental TOI and CBF values as well as the Fick-derived CMRO_2_. However, this optimization requires a demand decrease (Δp increase) and this causes a reduction in the Cu_A_ center. Now both the sign and magnitude of the modelled [oxCCO] differ from the experimental value. Modelling to optimize against all three experimentally measured parameters (Fig 6a) requires the increase in demand seen in the oxCCO only optimization. Now all modelled and experimental values show correct sign changes. However, the fit to the data is still rather poor, most clearly revealed by the model optimized CMRO_2_ again showing an increase in CMRO_2_. In short, modelling a CO_2_ effect on metabolic demand can *either* fit TOI, CBF and CMRO_2_ *or* [oxCCO], but not both. Primarily this conflict exists because a large *increase* in metabolic demand (uf) is required to explain the large increase in [oxCCO], whilst a small *reduction* in metabolic demand best fits TOI/CBF. Likewise, the parameter estimates for [R_autc_, τ_c_, τ_c2_, P_an_, CBF_n_] are unreliable because of the poor fit of the data (Table 1a).

In contrast, simulating changes in substrate supply (supp_s_) is reliable (Fig 7), illustrated by low error in measured versus modelled monitoring, and low standard deviation of parameter estimates (Table 1b). All three substrate supply optimizations (Fig 7) lead to a *decrease* in substrate supply, an increase in the NAD/NADH ratio, a fall in Δp and a fall in CMRO_2_. Optimizing against [oxCCO] alone (Fig. 7c) can model the measured increase in [oxCCO]. Although the absolute values of TOI and CBF are now modelled poorly, the modelled changes in these values are consistent with the magnitude of the experimental data. Therefore, in contrast with increases in demand (Fig. 6c), optimizing a change in substrate supply to better fit the [oxCCO] data alone *does* now agree with the Fick-derived fall in CMRO_2_. Optimizing to TOI/CBF alone (Fig 7b) with changes in substrate supply also provides acceptable agreement to all data sets (including interestingly both [oxCCO] and the Fick-derived CMRO_2_). Whilst additionally including [oxCCO] optimization (Fig. 7a) not surprisingly decreases the error in [oxCCO], this is at the cost of a poorer fit to CBF; however, CMRO2 still shows good agreement with the Fick value.

Some additional meaning can be attributed to parameter estimates either individually or together where the fit to the data is sufficient as in Fig. 7 and Table 1b. While “demand” (uf) and “substrate supply” (supp_s_) are the focus of this investigation, variation in other parameters are required to simulate the data. Consistent changes are seen for CO_2_ reactivity (R_autc_, increase), CO_2_ time constant (τ_c_ increase to 25sec) and autoregulation (P_an_, small increase) where there is an acceptable fit of the data in the supp_s_ simulations. These reflect changes within anticipated normal ranges for these parameters. The predicted delay for the metabolic effects of CO_2_ (τ_c2_) is prolonged several times compared to that of the vascular effects (τ_c_) in all cases. CBF_n_ a constant in the baseline status of the arterial resistance and vessel diameter varies significantly, but it is important to note that the effect of this taken together with the changes in P_an_ and R_autc_ is to maintain the absolute modelled CBF to a physiologically normal value during the baseline period of the challenge (0-100 sec, CBF panels of Figs 6 and 7).

To summarize, in hypercapnia although the TOI, CBF and Fick derived CMRO_2_ can be modelled by increases in CO_2_ decreasing metabolic demand *or* decreasing mitochondrial substrate supply, only the latter is also consistent with the observed changes in the NIRS [oxCCO] signal.

## 4. DISCUSSION

NIRS provides a non invasive measure of brain oxygenation and mitochondrial metabolism following changes in arterial carbon dioxide levels. When combined with measures of brain blood flow (such as TCD), brain CMRO_2_ can be calculated by the Fick principle, assuming the measured NIRS oxygenation (TOI) tracks venous oxygen saturation changes (Boas et al., 2003). It is also possible to measure CMRO_2_ changes by fitting systemic and NIRS-derived data to a dynamic systems model of brain blood flow and metabolism (Banaji et al., 2008). Our results showed that these two methodologies are complementary and provide novel insights into brain energy metabolism in the conscious adult human brain.

### 4.1 The effects of hypercapnia on cerebral metabolism

These were evaluated by three key physiological components: 1) oxygen extraction (TOI, TCD, SpO_2_), 2) mitochondrial oxidation ([oxCCO]) and 3) systemic physiology (CO_2_, SpO_2_, MABP). Using changes in systemic physiology alone the unoptimized BrainSignals model suggested a small (< 5%) increase in CMRO_2_ (Fig 2). Such a change is well within the variability seen in the literature which report anything from 35% increase (Horvath et al., 1994; Yang and Krasney, 1995; Jones et al., 2005) to a 30% decrease (Kliefoth et al., 1979; Zappe et al., 2008; Xu et al., 2011) in CMRO_2_ in anaesthetized mammals and awake humans. However, this small increase in CMRO_2_ was inconsistent with the 15% decrease in CMRO_2_ using the Fick equation and the measured changes in oxygen extraction (Fig 4); the large change in [oxCCO] seen in Fig 3 (> 0.5 micromolar increase in oxidation) also requires further explanation in our model. The challenge in optimizing the BrainSignals to our NIRS data was therefore to develop an optimized model consistent with *both* the decrease in CMRO_2_ seen in the oxygen extraction data (Fig 4) *and* the increase in oxidation at the level of cytochrome oxidase seen in the mitochondrial data (Fig 3).

There are several potential physiological mechanisms to perturb the mitochondrial respiratory chain and effect an increase in the oxidation state of mitochondrial cytochrome oxidase (Cooper et al., 1994; Banaji, 2006), some - but not all - of which are also consistent with a fall in CMRO_2_. Essentially these can be divided into three areas: an increase in the supply of oxidizing equivalents (oxygen), a decrease in the supply of reducing equivalents (carbohydrates/fats), or a change in the metabolic demand (via changes in the proton motive force). By optimizing the parameters of our BrainSignals model to the NIRS signals (both hemoglobin **and** mitochondrial), we were able to test which these possible mechanisms were consistent with the measured NIRS signals. These will be addressed in turn.

#### 4.1.1 Oxygen

An increase in pO_2_ consequent to an increase in CBF has the potential to oxidize cytochrome oxidase. However, the increase in [oxCCO] described in the present study is difficult to explain solely as a secondary consequence of CBF related increases in pO_2_. In the adult human brain, NIRS measured [oxCCO] increases are seen in both hypercapnia and hyperoxia (Tachtsidis et al., 2009; Kolyva et al., 2014). However, the increase in oxidation in hyperoxia is generally smaller (0.1 – 0.2 µM), compared to that seen in hypercapnia (0.2 - 0.5 µM) in the same subjects. The larger change in hypercapnia is difficult to explain in solely vascular terms. The unoptimized BrainSignals model includes both pO_2_ and pCO_2_ effects on CBF. The increase in CBF in hypercapnia will increase mitochondrial pO_2_ in the model and cause a small increase in the [oxCCO] signal (Fig 1). However, the magnitude of the modelled change is only 20% the measured change (Fig 3); any increase in mitochondrial pO_2_ caused by hypercapnia is therefore unlikely to be the major cause of the increase in [oxCCO] seen experimentally. The temporal characteristics of [oxCCO] and TOS also argue against pO_2_ changes as the sole explanatory factor. There is a substantial delay in [oxCCO] behind Vmca and TOS changes (τ_c2_ >180sec) suggesting a delayed metabolic effect rather than a direct pO_2_ association.

Animal studies support the conclusion that hypercapnia induced increases in [oxCCO] cannot be explained as a simple by product of changes in mitochondrial pO_2_. In the neonatal pig changes in [oxCCO] during manipulation of inspired oxygen and CO_2_ demonstrated that the hypercapnic [oxCCO] increase was insensitive to changes in hemoglobin oxygenation (Quaresima et al., 1998).

#### 4.1.2 Metabolic Demand

An increased oxidation of cytochrome oxidase could be driven by an increase in oxidative metabolism (Figs 5,6). Metabolic demand could increase the ADP/ATP ratio, decrease the proton motive force and oxidase cytochrome oxidase (Cooper et al., 1994; Banaji et al., 2008). However, a simulated increase in oxidative metabolism driving [oxCCO] changes, is a poor solution in our model as the predicted CMRO_2_ *increase* to account for the observed increase in [oxCCO] would need to be almost 30% (Fig 6). In agreement with this, even studies of functional activation, where an increase in CMRO_2_ can be assumed, demonstrate only small [oxCCO] increases - between 0.05 µM and 0.2 µM for visual and frontal cortical changes (Heekeren et al., 1999; Kolyva et al., 2012). Our modelling therefore provides no evidence to support an increase in mitochondrial functional demand in hypercapnia.

Animal data support only a minimal role for brain activation driven changes in mitochondrial ADP as a mechanism for altering mitochondrial redox state in hypercapnia. In neonatal dogs ^31^P MRS studies do indeed show a show a rise in the ADP/ATP ratio in hypercapnia. However, electrical activity drops (Yoshioka et al., 1995). In adult rats, ADP changes are equivocal in hypercapnia depending on which anesthetic is used (Litt et al., 1986). In spontaneously breathing animals hypercapnic effects on ^31^P MRS measures were limited to a decrease in intracellular pH, shown by the shift in the Pi peak, while the levels of ATP, phosphocreatine and Pi (and hence calculated ADP) were unchanged (Barrere et al., 1990).

#### 4.1.3 Substrate supply (reducing equivalents)

A decrease in the rate of substrate supply (reductant) to the mitochondrial electron transfer chain will increase the NAD/NADH ratio. In contrast to changes in demand, this decrease in supply can explain the large 0.58µM increase in [oxCCO] whilst remaining consistent with a small (5%) drop in CMRO_2_ predicted by the Fick calculation (Figs 5 and 7). A substrate supply limitation is also consistent with published animal models, which demonstrate an increase in the NAD/NADH following increases in CO_2_ (Gyulai et al., 1982).

An alteration in brain pH (Barrere et al., 1990) can potentially explain how an increase in pCO_2_ is able to cause such a decrease in the supply of mitochondrial reducing equivalents. In anaesthetized animal models pH can fall as low as 6.94 during hypercapnia, although it is unaltered during hypocapnia (Cady et al., 1987). A reduction in pH directly increases the oxidation state of cytochrome *c* and cytochrome oxidase in isolated mitochondria (Wilson et al., 1988) and purified enzyme systems (Thornstrom et al., 1988). This direct effect of mitochondrial acidification acting at the level of the enzyme might explain the oxidation of cytochrome oxidase previously seen in a newborn piglet model of hypercapnia (Quaresima et al., 1998; Springett et al., 2000).

However, we (Quaresima et al., 1998) and others (Folbergrova et al., 1975) have also suggested alternative metabolic explanations where a drop in pH acts indirectly on substrate supply to decrease CMRO_2_. For example, Siesjö and co-workers (Folbergrova et al., 1975) suggested a hypercania-induced reduction in pH could inhibit the glycolytic enzyme phosphofructokinase (PFK). In favor of this view was an hypercapnia-induced increase in the levels of brain glucose-6 phosphate and fructose-6 phosphate (the substrates for PFK), and a simultaneous fall in pyruvate and lactate (Folbergrova et al., 1975). This suggests a glycolytic control cross-over point somewhere between glucose-6 phosphate and pyruvate. Given that a reduction in pH is known to inhibit PFK - a key enzyme with the ability to limit glycolytic flux – it was argued that this was the most likely control point for the metabolic effects of hypercapnia (Folbergrova et al., 1975).

Consistent with glycolytic inhibition, Willie et al recently demonstrated hypercapnia-induced reduction in the cerebral metabolic rate for glucose and lactate in healthy human volunteers (Willie et al., 2015). These authors proposed that the observed efflux of glucose from the brain is consistent with an impairment of glycolysis. In contrast, Bain et al, whilst also demonstrating reduced CMRO_2_ (measured by arterial and jugular venous bulb blood gas measurements) consequent to hypercapnia in humans (Bain et al., 2016), could not demonstrate a decrease in non-oxidative metabolism; in fact there was a trend for non-oxidative carbohydrate metabolism to increase in prolonged hypercapnia.

In addition to the (somewhat) conflicting data regarding the impact on glycolysis of hypercapnia in adult humans, there are other limitations with the hypothesis that brain pH is the sole driver for the hypercapnia-induced increase in [oxCCO] seen in our study. One is its temporal profile; the [oxCCO] rise does not return rapidly to baseline. As the abrupt cessation of hypercapnia can be associated with cerebral alkalosis (Nioka et al., 1987), the persistent elevation of [oxCCO] is evidence against cerebral pH as the sole factor controlling [oxCCO] changes in hypercapnia.

Previous *in vitro* and *in vivo* studies also argue against a sole role for glycolysis as the target for CO_2_ effects on mitochondrial metabolism. In the same paper in which Siesjö and co-workers suggested that hypercapnia inhibited glycolysis, there was an intriguing additional effect on the TCA cycle intermediates (Folbergrova et al., 1975). Succinate levels increased, whereas those of fumarate decreased consistent with a control point for CO_2_ within the TCA cycle that interfaces directly with the electron transport chain at mitochondrial complex II (Folbergrova et al., 1975). This is consistent with both isolated mitochondrial studies that demonstrate inhibition by CO_2_ - bicarbonate mixtures of oxygen consumption at the level of succinate dehydrogenase (complex II) (Kasbekar, 1966) and studies on the purified succinate dehydrogenase enzyme that suggest two bicarbonate binding sites for inhibition, both with a Ki of 12mm i.e. well within the physiological range (Zeylemaker et al., 1970).

### 4.2 Effect of succinate dehydrogenase inhibition on cytochrome oxidase and the NIRS [oxCCO] signal

The mitochondrial electron transfer chain consists of four electron transfer complexes: complex I (NADH dehydrogenase), complex II (succinate dehydrogenase), complex III (the *bc*_1_ complex) and complex IV (cytochrome oxidase). The NIRS detectable Cu_A_ is the first redox center of complex IV, accepting the electrons from cytochrome *c* that will ultimately reduce oxygen to water. Cu_A_ is in redox equilibrium with cytochrome *c* (Cooper et al., 1997b; Mason et al., 2009) and will therefore track changes in cytochrome *c* caused by inhibitors of mitochondrial respiration (Chance et al., 1963).

Mitochondrial inhibitors acting upstream of cytochrome oxidase at complex I, II or III will slow the rate of reduction of Cu_A_, resulting in an oxidation that should effect an increase in the NIRS [oxCCO] signal (Chance et al., 1963; Cooper et al., 1997b; Banaji, 2006). On the other hand, complex IV inhibitors that act downstream of the Cu_A_ site and will reduce the Cu_A_ center and decrease the [oxCCO] signal, as is clearly seen with cyanide (Cooper et al., 1999). Interestingly it has been suggested that the vasodilator nitric oxide is in part responsible for the increase in blood flow seen in hypercapnia (Horvath et al., 1994). Nitric oxide is both a vasodilator and an inhibitor of mitochondrial respiration at the level of cytochrome oxidase (Cleeter et al., 1994). However, if in our studies NO was acting to decrease mitochondrial oxygen consumption we would expect a reduction of Cu_A_, not an oxidation (Brown and Cooper, 1994).

Succinate dehydrogenase is uniquely both a TCA cycle enzyme and a mitochondrial electron transfer complex (complex II). It therefore supplies electrons to complex I indirectly via the NADH production of the whole TCA cycle and to complex III directly via its own succinate dehydrogenase activity. Inhibiting this enzyme will therefore decrease the rate of electron supply to cytochrome oxidase by two mechanisms, leading to an inhibition of oxygen consumption, an oxidation of Cu_A_ and a consequent increase in the [oxCCO] signal (Banaji, 2006).

Therefore, a plausible mechanistic explanation of our measured and modelled data is that CO_2_ inhibition of succinate dehydrogenase activity in hypercapnia induces both a small CMRO_2_ decrease and an oxidation of mitochondrial cytochrome oxidase. This also has potential implications for the in vivo levels of succinate, a key intermediate in metabolism, free radical production, signal transduction, hypoxia, and tumorigenesis (Tretter et al., 2016).

### 4.3 Implications of hypercapnia inducing a fall in CMRO_2_

Whatever the molecular mechanism, the suggestion that CMRO_2_ may decrease during hypercapnia has significant implications. A hypercapnia-induced shift towards non-oxidative cerebral metabolism, and consequent decrease in CMRO_2_, may promote brain oxygen conservation and protect against severe apnea-related hypoxia (Bain et al., 2016). pCO_2_ is also a key in the manipulation of CBF and blood volume in the management of acute brain injured patients (Helmy et al., 2007; Stocchetti et al., 2017) and knowledge of its effect on CMRO_2_ is therefore clinically relevant. Finally, an incorrect assumption that CO_2_ is isometabolic would lead to underestimation of CMRO_2_ based on the CO_2_ challenge technique for BOLD calibration (Davis et al., 1998). Indeed mounting evidence of a metabolic effect of CO_2_ has stimulated investigation into alternative methods of MRI calibration (Peng et al., 2017) such as incorporating methods to increase CMRO_2_ and hence balance out its depressive effect.

### 4.4 Comparisons with other studies

Human studies using MRI and/or EEG have suggested no change (Chen and Pike, 2010; Jain et al., 2011) or a modest 8% (Peng et al., 2017) or 13% fall in CMRO_2_ during hypercapnia (Xu et al., 2011). Our NIRS and modelling work agrees with the latter studies. However, arguably all these datasets could be consistent given the likely small effect size, variation in measurement technique and small sample sizes (approximately 10 subjects in each). Alternatively, they may reflect temporal characteristics of the CO_2_ challenge. The studies used differing lengths of CO_2_ exposure; 180sec (Chen and Pike, 2010; Jain et al., 2011) and 360sec (Xu et al., 2011; Peng et al., 2017). Temporal resolution for these studies is also limited by the time taken for MRI resolution, which is generally of the order of 30sec (Jain et al., 2011).

In our study (300sec hypercapnia), the modelled prediction of a delayed onset of a metabolic effect (τ_c2_ >180sec) could theoretically explain this discrepancy, given that the two shortest (180sec) challenges were also the ones that showed no metabolic effect (Chen and Pike, 2010; Jain et al., 2011). We therefore conclude that all recent studies are consistent with our modelled small hypercapnia-induced decrease in CMRO_2_.

### 4.5 Limitations of this study

The Fick model for measuring CMRO_2_ using NIRS measured brain oxygenation as a surrogate for venous saturation is open to criticism. The calculation of CMRO_2_ using NIRS is sensitive to the accuracy of TOI, and assumes the arterial to venous ratio remains static; both of these factors are potentially problematic.

Absolute quantification of TOI is non-trivial given the degree of accuracy required for CMRO_2_ quantification. However, two key contributors to this accuracy are light attenuation due to water and the wavelength dependence of light scattering (Kleiser et al., 2016), both of which we have addressed via additional optical measurements.

NIRS offers high temporal resolution, but less spatial discrimination; TOI therefore reflects a combination of arterial, capillary and venous oxygenation, and thus OEF cannot be discriminated by focusing on specific vessels as in MRI. Physiological variability in the arterial venous ratio, particularly during the induction of CO_2_ has been demonstrated experimentally, and may violate the assumptions required for TOI and CBF to be used to directly calculate CMRO_2_ via a Fick equation (Moroz et al., 2012b). The method may therefore be poorly posed to discriminate between a small increase or decrease in CMRO_2_ (as noted for other methods above).

We have previously examined the behavior of TOI and the BrainSignals model following a hypercapnia challenge using a commercial NIRS device to measure TOI (Moroz et al., 2012b). TOI changes during hypercapnia were small and required a number of assumptions to be consistent with the observed CBF changes, most notably with regards to arterial/venous ratio changes, and extracerebral contamination of the signal. However, this requirement has largely been overcome in the present work by using an improved in-house broadband hybrid optical spectrometer and fitting to the group averaged data, rather than to each individual.

Robust CBF measurement is also required to calculate CMRO_2_ as TOI changes are interpreted in the context of estimated absolute or relative CBF. TCD of the middle cerebral artery measures changes in blood flow velocity rather than absolute cerebral blood flow; it is also targeted at a specific blood vessel, rather than the brain region covered by NIRS. Notwithstanding these differences relatively homogenous change in CBF would be anticipated with CO_2_ and thus the discrepancy between regional volumes is less relevant. The observed CBF reactivity of 4%/mmHg CO_2_ is also within normal expected limits, and model-based prediction of CBF from MABP and CO_2_ is successful. Cerebrovascular reactivity following CO_2_ administration is dependent on both vasodilation from CO_2_ and induced MABP changes – and it is therefore reassuring that the model replicates this behavior.

Our Fick-based hypothesis that CMRO_2_ fell during hypercapnia was further investigated using BrainSignals and 6 physiologically selected model parameters based on both our physiological questions of the data and parameter selection within other hypercapnia modelling studies (Ursino et al., 2000b; Moroz et al., 2012b). The BrainSignals model has been evaluated across a range of physiological paradigms in animal models (Caldwell et al., 2015), healthy volunteers (Jelfs et al., 2012) and brain injured patients (Highton et al., 2013). However as with all models of this form a degree of uncertainty exists in its replication of physiology.

An important advantage of combining broadband NIRS with a modelling approach such as BrainSignals over previous investigations which have only used a Fick model, is that it enables the measurement and interpretation of mitochondrial function, in this case the redox state of cytochrome oxidase via changes in NIRS measured [oxCCO]. Although there are multiple other parameters in our model that influence [oxCCO], as previously demonstrated in BrainSignals using sensitivity analysis (Caldwell et al., 2015), our primary aim here was to differentiate the effects of 3 key states: metabolic demand, pO_2_, and the effect of reductive substrate on [oxCCO] and CMRO_2_, all plausible and readily testable hypotheses previously discussed in the relevant literature.

The ability of NIRS to measure cytochrome oxidase redox state in the brain is potentially problematic as the concentration changes are generally much smaller than those seen in Hb (Matcher et al., 1995). However, [oxCCO] changes cannot readily be assigned to “crosstalk” from these larger changes (Uludag et al., 2004). Indeed in the case of hypercapnia, animal models showed that hypercapnia induced increases in [oxCCO] were identical when Hb changes were dramatically decreased following 80% replacement of blood by a perfluorocarbon blood substitute (Quaresima et al., 1998)

The [oxCCO] change is so large in our dataset that the only suitable explanation of the data with the current model and parameters is a reduction in substrate supply. The parameter predictions appear robust with a small standard deviation in repeated optimization evaluations. However, although this and previous (Tachtsidis et al., 2009; Kolyva et al., 2014) studies consistently show larger [oxCCO] changes in hypercapnia rather than hyperoxia, the absolute change in µM terms is less robust, relying as it does on terms that at present can only be approximated; these include both the total cytochrome oxidase Cu_A_ concentration that is NIRS detectable in the adult human brain and its resting redox state. In animal models the physiologically observed mitochondrial NIRS changes can be internally validated to a maximum possible [oxCCO] change in anoxia or following administration of mitochondrial inhibitors such as cyanide (Cooper et al., 1999), methods that are clearly not possible in human studies. Our suggestion that the mechanism of the CMRO_2_ fall is via inhibition of succinate dehydrogenase is therefore only tentative, depending as it does largely on the size of the cytochrome oxidase oxidation increase observed by NIRS. Indeed it has proven difficult, even in animal models, to increase [oxCCO] *in vivo* using inhibitors of complex I and II (Cooper et al., 1997a). Combination of ^31^P MRS (to evaluate ADP/ATP), NADH fluorescence and brain pO_2_ measurements combined with model based interpretation will still be required to shed further light on the intriguing effects of CO_2_ on brain energy metabolism.

## 5 CONCLUSION

Model assisted interpretation of cerebral [oxCCO] suggest that the increase in [oxCCO] following hypercapnia is consistent with a decrease in reductive substrate supply to the electron transport chain and a reduction in O_2_ consumption, consistent with the known bicarbonate inhibition of succinate dehydrogenase. This finding is supported by the emerging literature on the effect of hypercapnia on CMRO_2_ and glucose metabolism. The implications are far reaching because CO_2_ manipulation is frequently employed as a metabolically inert method of modifying CBF both clinically and in investigational techniques assessing CMRO_2_ and CBF. Further research is required in larger numbers of subjects to confirm our findings and might benefit from parallel animal investigation of hypercapnic changes in [ADP]/[ATP], mitochondrial pO_2_ and [NADH].

### Abbreviations

(Note abbreviations that are only used in Figures are described in the relevant figure Legend only).

*A*, NIR light attenuation; arterial pCO_2_ or PaCO_2_ , partial pressure of CO_2_ in arterial blood; BOLD, Blood Oxygen Level Dependent MRI; BrainSignals, a mathematical model of cerebral hemodynamics and metabolism; CBF, cerebral blood flow; CBFn, normal value of cerebral blood flow; CMRO_2_, cerebral metabolic rate for oxygen; Cu_A_, NIRS detectable binuclear Cu center in mitochondrial cytochrome oxidase; Dp model measured Δp; ETCO_2 ,_end-tidal CO_2_; θ, term representing the driving force for ATP synthesis in BrainSignals ; HOS, a hybrid NIRS optical system combining frequency domain and broadband (multiwavelength) measures of the NIR detectable light in tissue; L_CV_ the rate at which protons re-enter the mitochondrial matrix in BrainSignals controlled by the term *r*_CV_; MRI, Magnetic Resonance Imaging; fMRI functional Magnetic Resonance Imaging; MRS, Magnetic Resonance Spectroscopy; MABP, measured mean arterial blood pressure; NAD:NADH_rat_, mitochondrial NAD/NADH ratio; NIRS, Near Infrared Spectroscopy; oxCCO mitochondrial cytochrome oxidase redox state as measured by the oxidation of the NIRS-detectable Cu_A_ signal; Δ[oxCCO] change in concentration of oxidised Cu_A_ center; *ρ*, source detector separation in HOS; Δp, proton electrochemical potential comprising the mitochondrial membrane potential (Δψ) and the (ΔpH) across the inner mitochondrial membrane; P_an_, normal arterial blood pressure; PET, positron emission tomography; PFK, phosphofructokinase; R_autc_, magnitude of CO_2_ reactivity on vasculature; R_u_, control parameter in BrainSignals describing the direct effect of u on CBF; SpO2, hemoglobin saturation in arterial blood; supp, relative supply of reducing substrate to mitochondria; supps, influence of CO_2_ on supply of reducing substrates to mitochondria; τ_c_ time constant for vascular effects following changes in PaCO2; τ_c2_ time constant for metabolic effects following changes in PaCO2; TCA, tricarboxylic acid; TCD, transcranial Doppler; TOS, tissue oxygen hemoglobin saturation; TOI, Tissue Oxygenation Index - a NIRS measured approximation to TOS; *u*, a normalised term in the BrainSignals model encompassing brain metabolic demand (set to 1 in resting state normal brain); uf, influence of CO_2_ on *u*; Vmca, Blood velocity in the middle cerebral artery detectable by TCD; u^2DNADH^ term used in the BrainSignals model to indicate change in the rate of production of mitochondrial NADH via glycolysis and/or the citric acid cycle.

## Author Contributions

DH, MC, IT, CEE, MS, CEC designed the study. DH performed the research. DH, MC, CEC analysed the data. DH, MC, IT, CEE, MS, CEC wrote the paper.

## Supporting Information

**Fig S1**

Detailed description of BrainSignals Model

(1) Overview (2) Differential Equations, (3) Algebraic Equations (4) Chemical Reactions (5) State Variables (6) Intermediate Variables (7) Parameters (8) Derived Parameters

